# Observing non-covalent interactions in experimental electron density for macromolecular systems: A novel perspective for protein–ligand interaction research

**DOI:** 10.1101/2022.01.24.468575

**Authors:** Kang Ding, Shiqiu Yin, Zhongwei Li, Shiju Jiang, Yang Yang, Wenbiao Zhou, Yingsheng Zhang, Bo Huang

## Abstract

We report for the first time the use of experimental electron density (ED) in the Protein Data Bank for modeling non-covalent interactions (NCIs) for protein–ligand complexes. Our methodology is based on the reduced electron density gradient (RDG) theory describing intermolecular NCI by ED and its first derivative. We established a database called the Experimental NCI Database (ExptNCI; http://ncidatabase.stonewise.cn/#/nci) containing ED saddle points, indicating ~200,000 NCIs from over 12,000 protein–ligand complexes. We also demonstrated the use of the database for depicting amide–π interactions in a protein–ligand binding system. In summary, the database provides details on experimentally observed NCIs for protein–ligand complexes and can support future studies, including studies on rarely documented NCIs and the development of artificial intelligent models for protein–ligand binding prediction.

## INTRODUCTION

Non-covalent interactions (NCIs) govern protein–ligand interactions and are critical for understanding the determinants affecting ligand-binding affinity. To achieve a deep understanding of NCIs, many protein–ligand interaction databases have been established in the last decade.^1–7^ Two types of technologies are primarily applied to build such databases: (i) structure-based data mining and (ii) quantum mechanical (QM) methods-powered computation. For the first type, protein–ligand complex structures in the Protein Data Bank (PDB) are used as the main source, and different indices, such as distance, angle, exposed surface, and line-of-sight statistics, are used to depict the possibility of NCIs between a pair or two groups of atoms.^8–10^ For the second type, different levels of QM methods, ranging from semiempirical to coupled-cluster singles-doubles-and-triples wave function (CCSD(T)), are used to quantify the interaction energy of small model complexes.^5, 11^ The two technologies together have contributed greatly to the development of rules for the recognition of classical NCIs, such as hydrogen bonds, halogen bonds, salt bridges, and π-π stacking. To further expand the ability to recognize and quantify the entire spectrum of NCIs in highly complicated polarization environments such as protein–ligand binding systems and protein–protein interaction systems, we need to address the gap in the direct evidence of NCI between two proximal atoms in macromolecule systems. The gap is caused by the limitation of applying quantum mechanics for large systems and by the uncertainty of atom positions in the structures in PDB: e.g., the absence of hydrogen atoms and errors induced during structure building.

A potential solution for this gap can be found in the field of materials research^12^ in studies applying the reduced electron density gradient (RDG) theory^13^ in analyzing experimental electron density (ED) derived from the X-ray diffraction of small molecular crystals.^14, 15^ Stating the RDG theory in simple terms, NCI can be observed by pinpointing the ED saddle point, i.e. (3,−1) critical points and further quantified by measuring ED deviation from a homogeneous electron distribution using density and its first derivative (s = [1/(2(3π^2^)^1/3^)]|∇ρ|/ρ^4/3^). Some researchers have even proved that experimental ED can contribute to optimizing functions for density functional theory (DFT), given the fact that experimental ED is inherently time-averaged, whereas DFT ED represents pure ground-state.^12^

Inspired by research on small molecule crystals,^12, 14, 15^ we have developed a potentially path-breaking procedure to extract critical points from experimental ED for protein–ligand complexes deposited in the PDB. We processed >12,000 protein–ligand complexes and extracted ~200,000 saddle points. These data were subjected to noise reduction by varying the ED resolution and then consolidated into a database called the ExptNCI (Experimental NCI Database), available at http://ncidatabase.stonewise.cn/#/nci. In addition to database construction, we also present a case of using such data for empirical NCI mining. ED saddle points indicating amide–π interactions are extracted and used to support the QM interaction energy landscape scan. The QM result is well aligned with the observed points: 85% of the observed points are covered by the region with energy lower than −1.44 kcal/mol (semiempirical level). In addition to the attractive interaction of NH/π, which is consistent with previous research,^16^ we also found a −2.65 kcal/mol interaction (DFT level) between the edge of the aromatic ring and the amide plane when they interact in a perpendicular “edge-on” geometry.

## RESULTS

### NCI observed in experimental ED of the protein–ligand complex

X-ray diffraction (XRD) detects the electron distribution of the target molecule and generates an ED map. By searching for the ED saddle points, we can not only recognize classical NCIs, such as hydrogen bonds, π stacking, and halogen bonds, but also find relatively rare NCIs such as fluorine interacting with sulfur and methyl interacting with pyridine, as shown in Figure 1a–e. Additionally, because experimental ED represents a time-averaged density, some dynamics of the NCIs can also be observed (Figure 1f and Figure 2).

**Figure 1.**
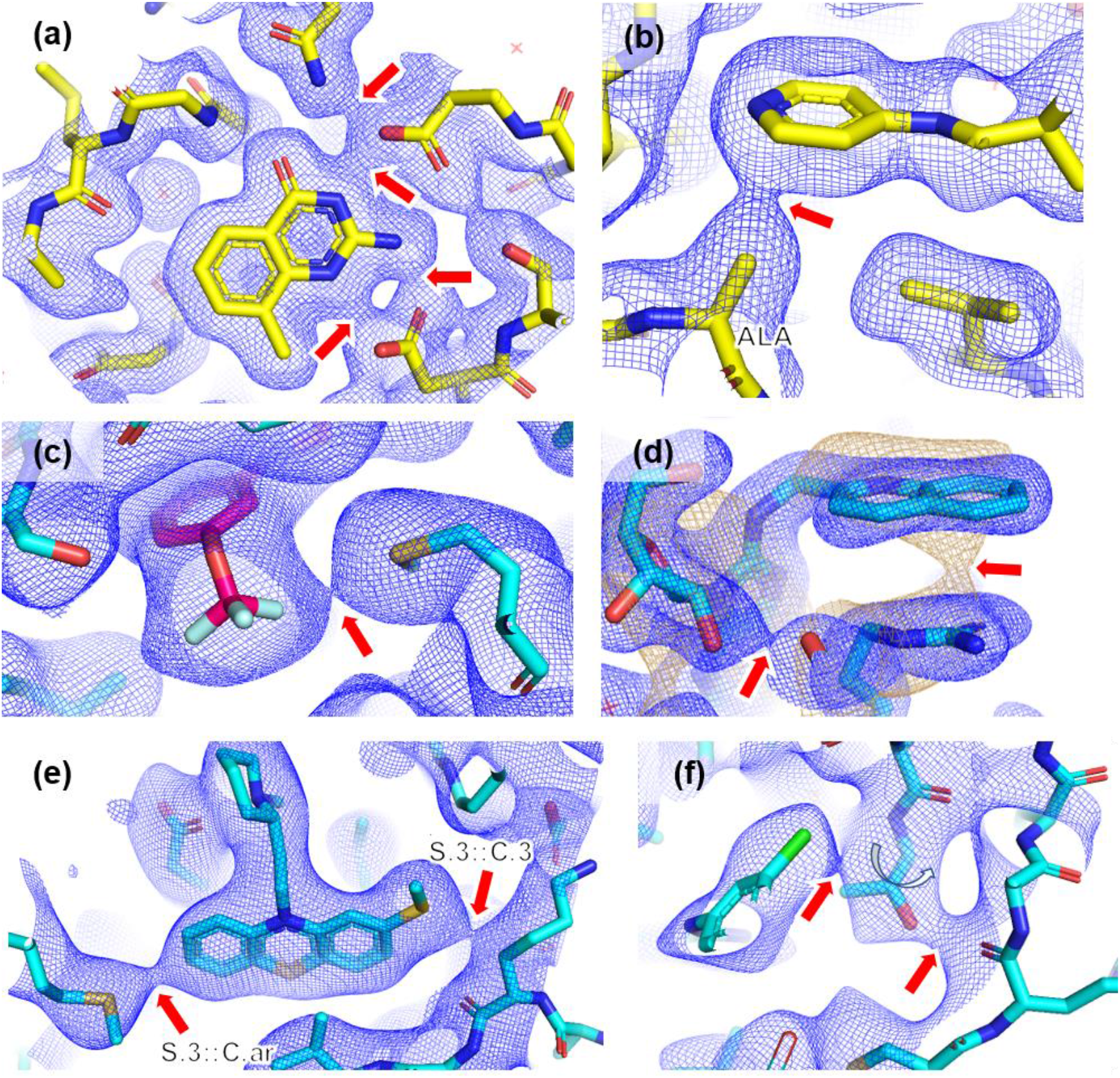
Observing NCI in X-ray diffraction-derived electron density maps (2Fo–Fc). Blue mesh indicates 2.5-Å resolution. All the maps are sigma-scaled and presented at specified counter levels. Saddle points are indicated by red arrows. a) Hydrogen bond interactions (PDB: 1S38, map counter level 0.2 sigma); b) Interaction between methyl and aromatic ring (PDB: 1Q8T, map counter level 0.2 sigma); c) interaction between F and methylthio (PDB: 2P4Y, map counter level 0 sigma); d) Weak π stacking revealed in low-resolution electron density map (PDB: 3LDQ; blue mesh indicates a 2.5-Å resolution map countered at 1.0 sigma; sand yellow mesh indicates a 3.5-Å resolution map countered at 1.0 sigma); e) Sulfur-involved NCI (PDB: 4I1R, map counter level 0.3 sigma); f) Observing NCIs under dynamic context caused by the rotation of threonine side chain (PDB:1XKK, map counter level 0 sigma).

**Figure 2.**
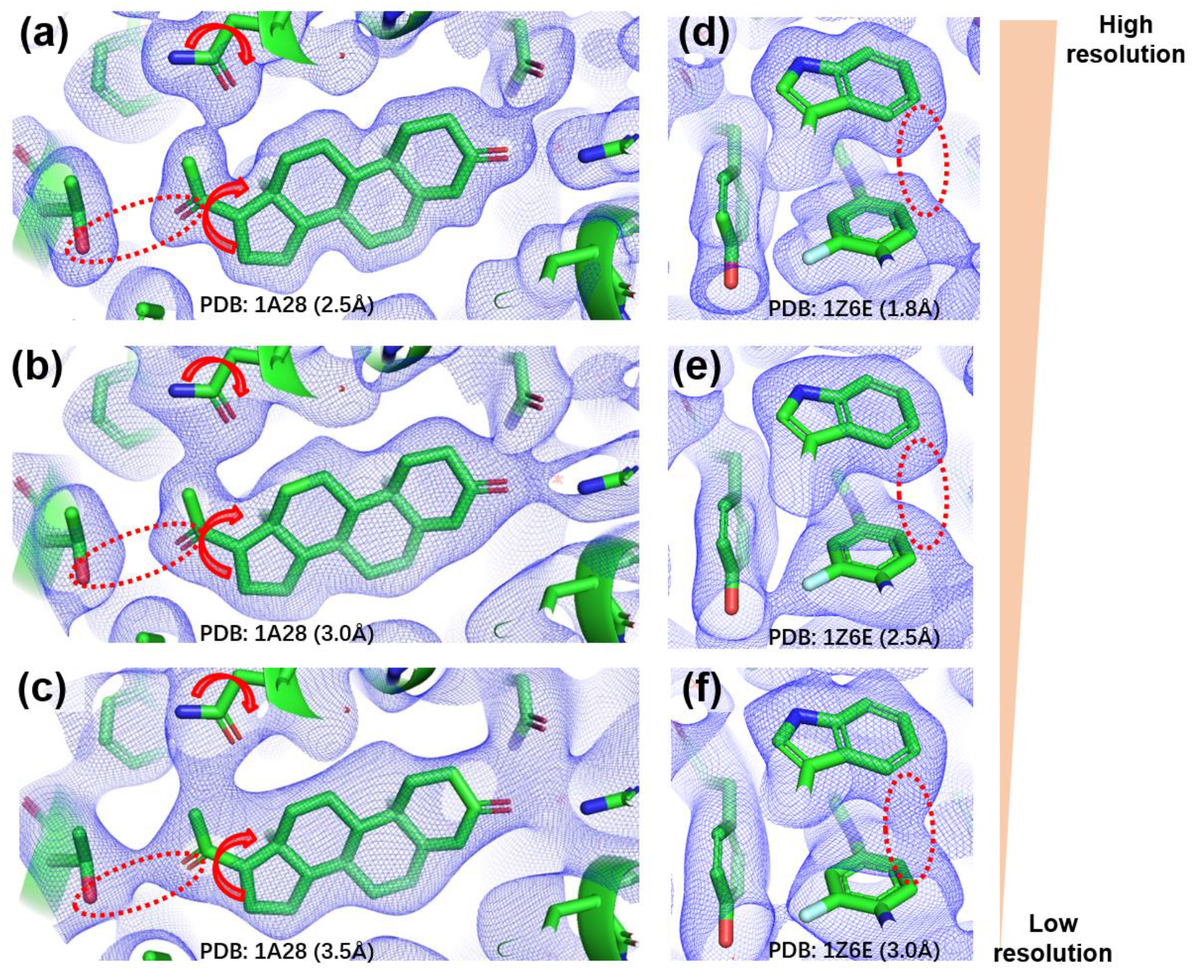
Emphasizing NCI signal in low-resolution ED maps. All the maps are sigma-scaled and presented at counter level 0 sigma. Red dashed circles indicate the relatively weak NCIs, which are clearly observed in low-resolution ED maps. Hydrogen bonds in a dynamic environment are shown in panels a, b, and c, with 2Fo-Fc maps for PDB 1A28 at 2.5-Å, 3.0-Å, and 3.5-Å resolution, respectively. Red arrows indicate the rotation of the groups causing the dynamics. π stacking contacts are shown in panels d, e, and f, with 2Fo-Fc maps for PDB 1Z6E at 1.8-Å, 2.5-Å, and 3.0-Å resolution, respectively.

Another benefit of using XRD ED for NCI detection is that we can clearly observe the signals of weak NCI; this can be done by checking them in low-resolution ED maps generated by only including XRDs at low-resolution. The intensity of XRD decreases as the ED map resolution increases, resulting in a relatively high signal-to-noise ratio for low-resolution ED maps, which enables us to confirm weak NCIs by checking them in ED maps at different resolutions (Figure 2). Doing so enables the identification of weak NCIs and helps distinguish them from false-positive signals. Our research conducted such a check in a low-resolution (3.5 Å) map for every NCI associated with saddle points having sigma-scaled intensity at 2.5 Å less than 0, i.e., less than the average.

In addition to using saddle points as a general indicator for recognizing NCI, we also used RDG in experimental ED as a more comprehensive NCI descriptor. Both repulsive and attractive interactions can be identified and visualized, as shown in Figure 3. Specifically, a spike in the RDG vs. sign(λ_2_)ρ plot indicates the presence of NCIs (Figure 3b), with the location of the spike on the negative side of the horizontal axis indicating an attractive interaction and that on the positive side of the axis indicating a repulsive interaction.^13^

**Figure 3.**
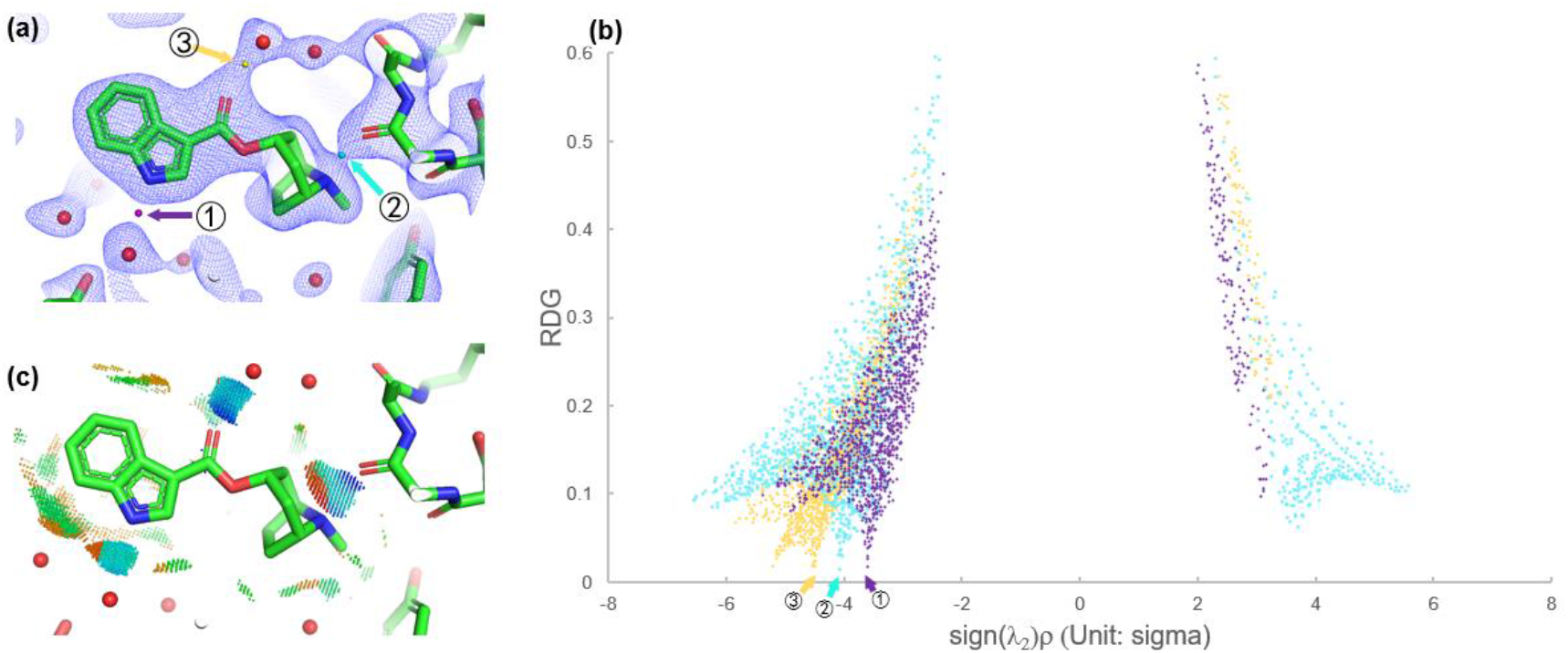
Depicting NCI with RDG in experimental ED for protein–ligand complex (PDB: 2WNC). a) Saddle points detected in 2fo–fc map (counter level 1.0 sigma). Three saddle points indicating three hydrogen bonds are respectively indicated by yellow, purple, and cyan arrows; b) plots of RDG versus electron density multiplied by the sign of the second Hessian eigenvalue for NCIs indicated in the panel (a) because the ρ here is sigma-scaled, to avoid negative value, all the ρ values used for calculating RDG and sign(λ_2_)ρ have their value added by 3. Spikes indicating three hydrogen bonds are indicated by arrows. All the dots on the scatter plot are colored according to their positions in real space. In detail, the dots within 1 Å of the saddle points 1, 2, and 3 are colored in yellow, purple, and cyan, respectively; c) RDG-based NCI isosurface showing the ligand–pocket interaction. Regions inside the RDG isosurface at a value of 0.2 (arbitrary unit) are indicated with dots, and the dots are colored based on sign(λ_2_)ρ using the rainbow scheme, in which blue depicts large negative values indicating strong, attractive interactions, and red depicts large positive values indicating repulsive interactions.

However, one limitation of using experimental ED for RDG analysis needs to be mentioned. Because of the lack of experimental measures on the forward-scattered reflection swamped by the transmitted beam, which is known as F_000_, the absolute value of ED is not available for macromolecule crystals. Therefore, the ED maps are contoured on a relative scale, and we had to use a sigma-scaled ρ for calculating RDG and sign(λ_2_)ρ. Consequently, the plot in Figure 3b has scales on the horizontal and vertical axes in arbitrary units. However, the spikes appearing in low-density regions still can indicate the occurrence of NCIs.

### ExptNCI database content

The current version of ExptNCI contains a total of 215,397 saddle points extracted from the experimental ED of 12,589 ligand–pocket complex structures in the PDB. The ED maps used for saddle points extraction have resolutions ranging from 2.5 to 4.5 Å, and 83% of them have a resolution greater than 2.5 Å. The ED topology information of the saddle points (such as sigma-scaled ρ, RDG, λ_1_, λ_2_, λ_3_, and Laplacian) as well as the structural information of atoms at both ends of the saddle points (such as residual name, element, and its hybridization in the Mol2/Sybyl atom format)^17^ are included in the database (Table 1). We also included ρ at a low-resolution (3.5 Å) at the position of saddle points in a 2.5-Å ED map to use it to distinguish noise from signals of weak NCIs. As discussed in the first part of the results section, blurring the map by only including low-resolution data with a relatively high signal-to-noise ratio can present weak NCIs more clearly. Here, we filtered out false-positive saddle points in a 2.5-Å resolution ED map to check if such points have negative sigma ρ in a 3.5-Å resolution ED map.

**Table 1.** List of fields in ExptNCI database

| Field | Description |
| --- | --- |
| NCI ID | NCI ID in ExptNCI Database |
| Complex ID | Complex ID composed by PDBID and ligand name |
| PDB Code | PDB Code |
| L_type | Ligand atom type in Mol2/Sybyl format |
| R_type | Receptor atom type in Mol2/Sybyl format |
| NCI_atom_pair | NCI classified based on Mol2/Sybyl formatted type of atoms at both ends of the saddle point |
| NCI_intuitive | NCI classified based on the properties of the atoms at both ends of the saddle points |
| NCI_ODDT | ODDT based NCI type (hydrogen bond, salt bridge, halogen bond, pi stacking, pi cation) |
| Resolution | The high-resolution limit of electron density map available in PDB |
| CP_type | Type of critical point: (3,-1) indicates saddle points while (3,+1) indicates ring CPs |
| RDG | $[1/(2(3\pi^2)^{1/3})]\vert \nabla\rho\vert /\rho^{4/3}$, where $\rho$ indicates the modified intensity of electron density. RDG is used as an indicator for NCI. |
| slr | $\text{sign}(\lambda_2)\rho$, where $\rho$ indicates the modified intensity of electron density at the critical point; $\lambda_2$ indicates the second larger value of the three eigenvalues of the electron density Hessian (second derivative) matrix. The value of slr is $\rho$ when $\lambda_2$ is positive, indicating repulsive interaction; it is $-\rho$ when $\lambda_2$ is negative, indicating attractive interaction. |
| $\lambda_1$ | Three eigenvalues of the electron density Hessian (second derivative) matrix, such that ($\lambda_1 \leq \lambda_2 \leq \lambda_3$) |
| $\lambda_2$ | Three eigenvalues of the electron density Hessian (second derivative) matrix, such that ($\lambda_1 \leq \lambda_2 \leq \lambda_3$), a positive $\lambda_2$ indicates repulsive interaction, while a negative $\lambda_2$ indicates attractive interaction |
| $\lambda_3$ | Three eigenvalues of the electron density Hessian (second derivative) matrix, such that ($\lambda_1 \leq \lambda_2 \leq \lambda_3$) |
| Laplacian | $\nabla^2\rho = \lambda_1 + \lambda_2 + \lambda_3$ |
| ED_2.5A_modified | $\rho + 3$, where $r$ is sigma-scaled ED intensity of the saddle point in 2Fo-Fc map at 2.5 Å. Such modified $\rho$ is used to calculate $\text{sign}(\lambda_2)\rho$ and RDG. |
| ED_2.5A | Sigma-scaled ED intensity of the saddle point in 2Fo-Fc map at 2.5 Å |
| ED_3.5A | Sigma-scaled ED intensity of the saddle point in 2Fo-Fc map at 3.5 Å |
| Distance | Distance between ligand atom and receptor atom |
| is_backbone | Boolean: 1 if NCI occurs on protein backbone |
| is_ED_based | Boolean: 1 if NCI is defined based on an ED saddle point |
| is_ODDT | Boolean: 1 if NCI is recognized by ODDT |
| ResName | Name of the receptor residue involved in the NCI |
| ResAtomName | Name of the receptor atom involved in the NCI |
| ResID | Residue ID of the receptor residue involved in the NCI |
| ChainID | Chain ID of the receptor residue involved in the NCI |
| LigName | Name of the ligand involved in the NCI |
| LigAtomName | Name of the ligand atom involved in the NCI |
| Lig_ResID | Residue ID of the ligand residue involved in the NCI |
| Lig_ChainID | Chain ID of the ligand residue involved in the NCI |
| Group | Type of atom in terms of protein, water, or heteroatoms |

By including strong saddle points with sigma-scaled intensity at 2.5-Å resolution above 0 and weak saddle points which can pass the false-positive test by using the low-resolution map of 3.5 Å (the method described above), we selected 95,532 saddle points, accounting for 51% of the originally labeled points (Figure 4a). Among them, 32% were also recognized as NCIs by rules embedded in the widely used software ODDT^18^, with hydrogen bonds accounting for the majority (Figure 4b). For the 68% that were not recognized by ODDT, we made a rough classification based on the properties of the atoms at both ends of the saddle points, as shown in Figure 1c, in which polar interactions (hydrophilic–hydrophilic), aliphatic C…hydrophilic (N/O) interactions, and aromatic…hydrophilic (N/O) interactions accounted for the majority.

**Figure 4.**
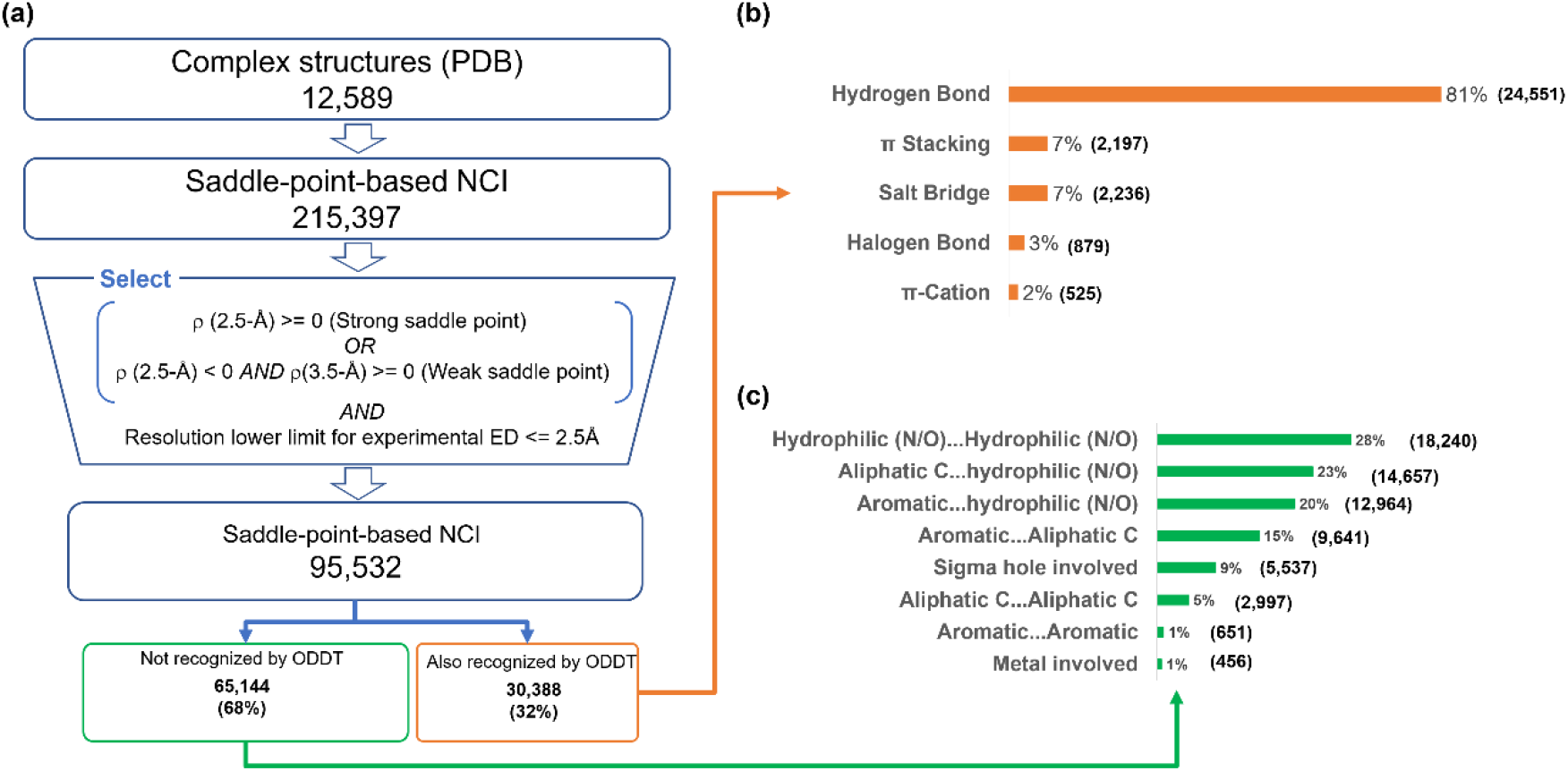
Database construction and dataset profile. a) Database construction workflow. *AND* and *OR* in the select box are logical operators; b) distribution of interaction type for NCIs recognized by both ODDT and ED saddle points; c) distribution of interaction type for NCIs recognized by ED saddle points but not ODDT.

### Usage Case: Depicting amide–π interactions in the ligand–protein binding system

Amide–π interactions^16^ have been increasingly studied for their involvement in the binding of drug molecules to target proteins.^19–21^ Most previous studies focused on how the plane of the arene ring interacts with the amide,^16, 20–22^ which can be classified as focusing on face-on geometry, a configuration with γ approximately 0° in a coordinate system, as shown in Figure 5a. To check whether such face-on geometry represents the majority of amide–π interactions in the protein–ligand binding system, we extracted 3,162 amide-π pairs from the ExptNCI database (details of the list provided in supplementary information). The amide–π pairs were extracted based on the fulfillment of the following requirements: (i) it must have ED saddle points between the aromatic carbon and any atom of the amide group and (ii) the ED map must have a resolution better than 2.5 Å (examples shown in Figure 5b). Those with saddle points between C=O and hetero atoms in the aromatic ring were excluded so that classical hydrogen bonds are not included in the analysis. The spatial distribution of the aromatic ring center relative to the carbon atom of the amide plane was plotted and colored with a γ-related color scheme. Interestingly, most of the interactions displayed γ values of approximately 90°, indicating an edge-on geometry (Figure 5c). Notably, face-on and edge-on interactions occur on two ellipsoids with different radii (Figure 5d and 5e).

**Figure 5.**
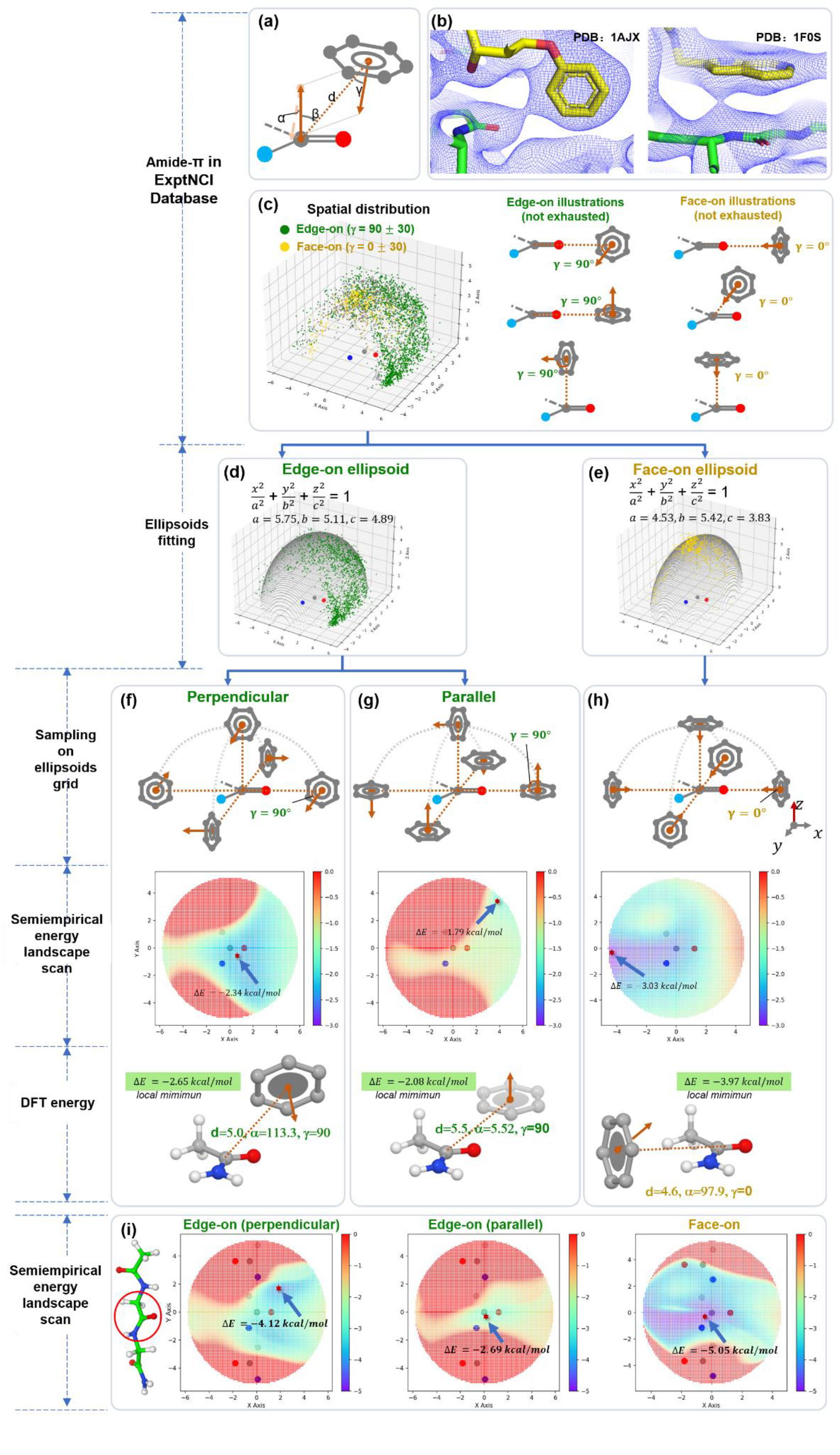
Using experimental ED data to support the profiling of amide–π interaction. a) Examples of amide–π interaction identified by ED saddle points. 2fo–fc map is countered at 0.3 sigma; b) coordinate system of amide–π interaction; c) spatial distribution of aromatic ring center relative to the carbon atom of the amide plane. Green and yellow indicate edge-on and face-on geometry, respectively. Illustrations of edge-on and face-on are also provided; (d) and (e) are ellipsoids and parameters obtained by fitting edge-on and face-on positions, respectively, to the general equation of an ellipsoid; for (f), (g), and (h), the top part represents the sampling scheme of formamide– benzene conformation on the ellipsoids, with edge-on conformation sampled in two ways: perpendicular and parallel; the middle part represents GFN2-xTB level energy landscape, with the blue arrow pointing to a red star indicating global minimum; bottom part represents the conformation for global minimum on GFN2-xTB energy landscape and its M06-2x/6-311+G(d,p) energy calculated by GAMESS; i) energy landscape scan for N-acetyl glycyl glycinamide. The ellipsoid is with respect to the amide group indicated by the red circle.

To further investigate the interaction geometry for amide–π, we identified the ellipsoids for face-on and edge-on geometry by fitting the aromatic center positions of the two types of geometry to the general equation of an ellipsoid (Figure 5d and 5e). Subsequently, we computed the GFN2-xTB level energy landscape based on the fitted ellipsoids using a formamide–benzene model system (Figure 5f, 5g, and 5h). For edge-on geometry (i.e., γ=90), the interaction is favored when benzene approaches the amide plane from the top of C=O perpendicularly (Figure 5f and 5g), with a minimum interaction energy of −2.65 kcal/mol calculated using M06-2x/6-311+G(d,p). For face-on geometry (i.e., γ=0), the result of our energy landscape scan is consistent with previous studies,^16^ showing a favored interaction of NH/π and a repulsive interaction of C=O/π, as shown in Figure 5h. The same approach was also applied to the amide group in a tripeptide to simulate the situation in the protein (Figure 5i). The computed energy landscape enjoyed a decent match to the spatial distribution of the observed amide–π interactions extracted from ExptNCI, with 85% of the latter covered by the former region with energy lower than −1.44 kcal/mol.

In summary, the use of observed ED saddle points for NCI description is demonstrated in this case through its support for an energy landscape scan.

## DISCUSSION

XRD provides an experimental ED map that contains massive amounts of information. Partial information is effectively interpreted into atom coordinates, and this information is entered in the PDB. However, in addition to atom coordinates, there is still plenty of information hidden in the experimental ED maps. For the first time, we extracted NCI signals from the ED maps and used them to establish the ExptNCI database.

How does the ED-saddle-point-based observation complement and further improve geometry-rule-based description? In most cases, the rule-based NCI descriptions have the following two characteristics: 1) they focus on a pair of atoms or groups by simplifying the environment and 2) they highly rely on the precision of atomic coordinates. Therefore, such description cannot always appropriately profile the NCIs in practical situations where the polarization environment is compilated and the model structures are inaccurate or have missing regions. Because ED-saddle-point-based NCI observation mainly depends on electron density, it is less sensitive to the accuracy of the coordinates than rule-based descriptions. In addition, because the saddle points are detected from the experimental electron density, they can reflect the complex polarization environment and provide more information such as the relative strength of the NCIs and geometrically rare cases to support the development of new empirical rules. These superiorities are shown in two cases (Fig. S1 and S2), where the saddle-point-based method is compared with that of the rules-based method for sulfur-involved NCIs and π-π stacking.^18, 23, 24^

When exploring the ExptNCI database, users should check the three following aspects if some seemingly unusual NCIs are found: (i) check whether the structure is correctly determined, which can be judged by checking positive or negative densities around the NCI region of interest in the Fo–Fc map; (ii) check whether low-resolution causes merging of saddle points. An ED map becomes less detailed when the resolution is low, and two proximal saddle points may merge into one in a low-resolution ED map. As shown in Figure 6, just because there is only one saddle point between C=O and C=O in a 2.7-Å resolution ED map, it does not necessarily indicate the existence of NCIs between the two *sp^2^* oxygen atoms. In other words, the case in Figure 6 resulted from the merging of two saddle points representing two individual classical hydrogen bonds; (iii) check whether there are any dynamics that can make the interaction more reasonable, e.g., the flip of the side chain for Gln, Asn, and His.

**Figure 6.**
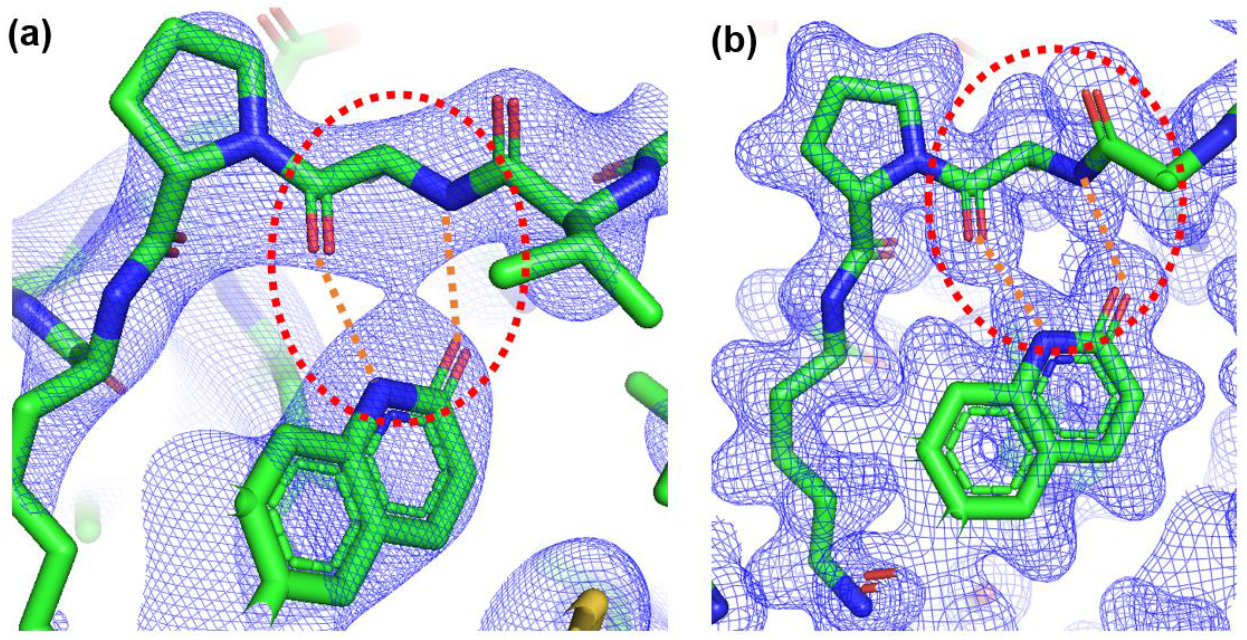
Merging saddle points in low-resolution ED maps (PDB: 6MA1). a) Experimental ED 2Fo–Fc map at 2.7-Å resolution shown at the counter level of 1.4 sigma. Two classical hydrogen bonds, indicated by orange dash lines, exist within the red dashed circle, but only one saddle point is observed; b) GFN2-xTB calculated electron density, showing two saddle points at the same region. The map is countered at 0.03 e^−^/Å^3^. The empirical QM calculation is conducted using xtb^25^.

To further improve the data in terms of quantity and quality, we consider two directions. The first is to expand the scale of the database by extracting NCIs from the interface of protein–protein interactions (PPI). This may allow us to achieve a more detailed understanding of the interaction fingerprint and ultimately benefit peptide/protein design. The second direction is to improve the accuracy of the data by solving multi-crystal variance, which is a problem caused by the lack of absolute ED values for macromolecule crystals. Such a challenge could be tackled by converting the ED values from the sigma-scaled density to the number of electrons. Previous studies that were aimed at measuring the quality of structures in PDB by analyzing ED can serve as a good starting point.^26, 27^

Including the experimental saddle point ED intensity as NCI information can also be considered as a solution to support artificial intelligence-based protein–ligand binding prediction. Although experimental NCI is not always available as input, because most often, pocket–ligand complexes are generated by docking or molecular dynamics and thus lack experimental ED, we can build two machine learning models to first predict NCI from a given protein–ligand complex structure, and then use the predicted NCI to facilitate ligand binding affinity prediction.

In addition to providing more data resources, describing NCI from the perspective of crystallography ED also inspired us to consider leveraging crystallography as a solution for molecular representation for machine learning models. To date, the majority of attempts by researchers to find molecular representations have been in real space, and many reports have been made using strings, molecular graphs, molecular matrixes, potential fields, and atom density fields.^28^ However, an ideal representation comprehensively reflecting physical and chemical information, friendly to mathematics, and supported with plenty of experimental data available for AI model training is still absent. By applying crystallography theory, we can further expand the attempt in reciprocal space (i.e., frequency domain) and take a big step forward to realizing the ideal representation for molecules. Specifically, we apply Fourier transform (FT) on the atomic coordinates to transfer the information from real space to the frequency domain and then apply reverse FT on the frequency domain to bring back the information to real space as ED. By varying the resolution when conducting reverse FT in the frequency domain, we can obtain ED in real space with different levels of detail, emphasizing scaffold, atom, or even bond properties. Unlike graphs composed of vertices and edges, such representations fill the space in a continuously differentiable manner, which is favored by the CNN model. Unlike other 3D molecular representations, such representations are naturally associated with a large amount of testing data: the experimental ED deposited in the PDB. We have already tested them on a 3D molecule generation model and have obtained some promising results that will be reported later.

In summary, a massive amount of information is present in the experimental ED maps deposited in the PDB. The usage of only part of that information has created our current understanding of protein structures. We hope that our work can shed some light on leveraging experimental ED maps to further understand NCIs in the macromolecular system and on combining crystallography and AI from the perspective of providing reliable data sources and exploring better representation of molecules.

## METHODS

### Database construction

#### Experimental ED map processing and critical point labeling

All coordinates and map coefficients were obtained from PDB-REDO.^29^ ED maps covering ligands and pocket residues within 5 Å of the ligands were synthesized at multiple resolutions using Phenix.^30^ The maps were stored in the xplor format with a 0.15-Å grid interval. The critical points were labeled using the following procedure:

The ligand in our database is defined using PDBbind (version 2019) as a benchmark.^31^ Ligand/receptor atom pairs with a distance <5 Å were identified, and the midpoint was set as the origin;

The RDG value of all the grids was calculated within 1 Å of the origin;

The gird point with local minimum RDG was found and marked as a saddle point candidate;

For all the saddle point candidates, the eigenvalue of the Hessian matrix was calculated and sorted so that λ_3_> λ_2_>λ_1_. If the eigenvalues did not fulfill the criteria of λ_3_> λ_2_> λ_1_, the candidate was discarded;

If there were two saddle point candidates <0.5 Å from each other, the one with the relatively weaker intensity was discarded.

#### Atom property annotation

The topology of ligands from PDB entries was curated by RDKit with isosteric SMILES from RCSB Ligand-Expo, and other ligands with missing data were curated using OpenBabel. The Mol2/Sybyl atom types of pockets and ligands in the database were annotated using OpenBabel and PyBel packages, and the rule-based molecular interactions in the database were analyzed and classified using the ODDT software package (version 0.7).^18^

#### Web interface implementation

The database website was developed with a Java backend. The ligand similarity search or substructure search in the database was developed using RDKit, and NCI information was stored and queried through MySQL. NGL.js was implemented to display the receptor–ligand complex and the ED map.

#### Amide–π interaction model

To avoid including O..N.ar hydrogen bonds, only amide–π systems with ED saddle points between C/N/O on the protein backbones and C.ar on ligands were subject to our analysis. To profile the spatial distribution of aromatic ring centers, all the amide groups of interest were superimposed and placed on the X––Y plane with a uniform orientation (Figure 5b), and all the aromatic centers of the amide–π systems were plotted in the Z-positive sector, given that the amide plane is a mirror plane.

Four parameters including angles α, β, γ, and distance d are defined as shown in Figure 5b to describe amide–π geometry, in which the angle α is used to describe whether the π system is parallel (α=0 °±30° or 180 °±30°) or perpendicular (α=90±30°) to the amide plane; the angle γ is used to describe whether the aromatic ring center is facing toward the amide group in a “face-on” geometry (γ=0±30°), or showing its edge toward the amide group in an “edge-on” geometry (γ=90±30°).

Ellipsoids for face-on and edge-on geometry were identified by fitting the aromatic center position for the two types of geometry to the general equation of an ellipsoid (Figure 5d, 5e). Then the fitted ellipsoids are represented by grids with an interval of 0.1 Å along both X and Y axes. To scan the interaction energy landscape based on fitted ellipsoids for benzene and formamide systems, we first determined the zero-point energy (−29.70 kcal/mol) by applying GFN2-xTB calculation on a benzene–formamide complex with a distance of 50 Å between the two groups. Afterward, we placed benzene on the face-on grid in the pose where γ equals 0° to obtain a face-on geometry complex subset (Figure 5h). For the edge-on geometry complex subset, when we placed a benzene group on the grid of edge-on ellipsoids, there were two types of poses that fulfilled the requirement of γ=90°. Therefore, we divided the edge-on geometry into perpendicular-edge-on and parallel-edge-on subtypes. In detail, we used a plane defined by the Z-axis and the vector connecting the amide carbon to the benzene center to distinguish the two subtypes: if the norm of benzene was in the above-defined plane, then it was a parallel-edge-on sub-type (Figure 5f); if the norm of benzene was perpendicular to the above-defined plane, then it was a perpendicular-edge-on sub-type (Figure 5g). The energy landscapes for the geometries of face-on, parallel-edge-on, and perpendicular-edge-on were synthesized by calculating GFN2-xTB energy and then subtracting the zero-point energy from it for the complexes on the corresponding grids. Complexes representing the global minimum of the three energy landscapes were also subjected to the M06-2x/6-311+G(d,p) calculation using GAMESS to obtain the DFT level energy. The energy landscape scan for benzene interacting with amide groups in the context of tripeptide was conducted in a similar way using a GFN2-xTB-optimized N-acetyl glycyl glycinamide as a starting point. Because the optimized molecule is not subject to mirror symmetry, we scanned the entire ellipsoid and combined the upper and lower halves by overlapping the grids of the two parts and using the lower energy on the two overlapped grids as the final value to compose the energy landscape.

#### Software for figures and tables

The structure and ED figures were made using Pymol.^32^ Statistical analysis was performed using Pandas^33^ and Numpy packages.^34^ Scatter plots were constructed using Matplotlib^35^ and Inkscape.^36^

## ABBREVIATIONS

ED: electron density
NCI: non-covalent interaction
PDB: Protein Data Bank
RDG: reduced electron density gradient
QM: quantum mechanical
DFT: density functional theory
XRD: X-ray diffraction
PPI: protein–protein interaction
FT: Fourier transform

## Author Contributions

B. H. and Y. Z. conceived the idea. B.H. provided instructions for all experiments. W. Z. provided instructions on AI models. K. D. constructed the database. S. Y. developed the saddle point labeling script. Z. L. supported the saddle point labeling and performed quantum mechanics calculations. S. J. implemented the web interface for the database. Y. Y. designed the web interface.

## Funding Sources

All the authors received funding from StoneWise Ltd. B.H. also received funding for cloud computing from Beijing Municipal Science & Technology Commission project Z211100003521001.

## Data availability

http://ncidatabase.stonewise.cn/#/nci

## Code availability

Available upon request.

## Competing Interests

The authors declare no competing interests.

## ACKNOWLEDGMENT

## Notes

http://ncidatabase.stonewise.cn/#/nci

### Competing Interest Statement

The authors have declared no competing interest.

### Summary of Updates

A paragraph is added in the discussion sector to further clarify how the ED-saddle-point-based observation can complement and further improve the geometry-rule-based description

## REFERENCES

1. Anand, P.; Nagarajan, D.; Mukherjee, S.; Chandra, N., PLIC: Protein-Ligand Interaction Clusters. Database (Oxford) 2014, 2014, bau029.

2. Angles, R.; Arenas-Salinas, M.; García, R.; Reyes-Suarez, J. A.; Pohl, E., GSP4PDB: A Web Tool To Visualize, Search And Explore Protein-Ligand Structural Patterns. BMC Bioinformatics 2020, 21, 85.

3. Gallina, A. M.; Bisignano, P.; Bergamino, M.; Bordo, D., PLI: A Web-Based Tool For The Comparison of Protein-Ligand Interactions Observed on PDB Structures. Bioinformatics 2012, 29, 395–397.

4. Inhester, T.; Rarey, M., Protein–ligand Interaction Databases: Advanced Tools To Mine Activity Data And Interactions On A Structural Level. 2014, 4, 562–575.

5. Jurecka, P.; Sponer, J.; Cerný, J.; Hobza, P., Benchmark Database Of Accurate (MP2 And CCSD(T) Complete Basis Set Limit) Interaction Energies Of Small Model Complexes, DNA Base Pairs, And Amino Acid Pairs. Physical chemistry chemical physics: PCCP 2006, 8, 1985–93.

6. Murakami, Y.; Omori, S.; Kinoshita, K., NLDB: A Database For 3D Protein-Ligand Interactions In Enzymatic Reactions. Journal of structural and functional genomics 2016, 17, 101–110.

7. Rezac, J., Non-Covalent Interactions Atlas Benchmark Data Sets: Hydrogen Bonding. J Chem Theory Comput 2020, 16, 2355–2368.

8. Ferreira de Freitas, R.; Schapira, M., A Systematic Analysis Of Atomic Protein-Ligand Interactions In The PDB. Medchemcomm 2017, 8, 1970–1981.

9. Kuhn, B.; Gilberg, E.; Taylor, R.; Cole, J.; Korb, O., How Significant Are Unusual Protein-Ligand Interactions? Insights from Database Mining. J Med Chem 2019, 62, 10441–10455.

10. Xu, Z.; Zhang, Q.; Shi, J.; Zhu, W., Underestimated Noncovalent Interactions in Protein Data Bank. J Chem Inf Model 2019, 59, 3389–3399.

11. Hobza, P., Calculations On Non-Covalent Interactions And Databases Of Benchmark Interaction Energies. Acc Chem Res 2012, 45, 663–72.

12. Kasai, H.; Tolborg, K.; Sist, M.; Zhang, J.; Hathwar, V. R.; Filso, M. O.; Cenedese, S.; Sugimoto, K.; Overgaard, J.; Nishibori, E.; Iversen, B. B., X-Ray Electron Density Investigation Of Chemical Bonding In Van Der Waals Materials. Nat Mater 2018, 17, 249–252.

13. Johnson, E. R.; Keinan, S.; Mori-Sanchez, P.; Contreras-Garcia, J.; Cohen, A. J.; Yang, W., Revealing Non-Covalent Interactions. J Am Chem Soc 2010, 132, 6498–506.

14. Saleh, G.; Gatti, C.; Lo Presti, L., Non-Covalent Interaction Via The Reduced Density Gradient: Independent Atom Model Vs. Experimental Multipolar Electron Densities. Computational and Theoretical Chemistry 2012, 998, 148–163.

15. Saleh, G.; Gatti, C.; Lo Presti, L.; Contreras-Garcia, J., Revealing Non-Covalent Interactions In Molecular Crystals Through Their Experimental Electron Densities. Chemistry 2012, 18, 15523–36.

16. Imai, Y. N.; Inoue, Y.; Nakanishi, I.; Kitaura, K., Amide-Pi Interactions Between Formamide And Benzene. J Comput Chem 2009, 30, 2267–76.

17. Clark, M.; Cramer III, R. D.; Van Opdenbosch, N., Validation Of The General Purpose Tripos 5.2 Force Field. 1989, 10, 982–1012.

18. Wojcikowski, M.; Zielenkiewicz, P.; Siedlecki, P., Open Drug Discovery Toolkit (ODDT): A New Open-Source Player In The Drug Discovery Field. J Cheminform 2015, 7, 26.

19. Krone, M. W.; Travis, C. R.; Lee, G. Y.; Eckvahl, H. J.; Houk, K. N.; Waters, M. L., More Than pi-pi-pi Stacking: Contribution of Amide-pi and CH-pi Interactions to Crotonyllysine Binding by the AF9 YEATS Domain. J Am Chem Soc 2020, 142, 17048–17056.

20. DeFrees, K.; Kemp, M. T.; ElHilali-Pollard, X.; Zhang, X.; Mohamed, A.; Chen, Y.; Renslo, A. R., An Empirical Study of Amide-Heteroarene pi-Stacking Interactions Using Reversible Inhibitors of a Bacterial Serine Hydrolase. Org Chem Front 2019, 6, 1749–1756.

21. Bootsma, A. N.; Wheeler, S. E., Stacking Interactions of Heterocyclic Drug Fragments with Protein Amide Backbones. ChemMedChem 2018, 13, 835–841.

22. Harder, M.; Kuhn, B.; Diederich, F., Efficient Stacking On Protein Amide Fragments. ChemMedChem 2013, 8, 397–404.

23. Koebel, M. R.; Cooper, A.; Schmadeke, G.; Jeon, S.; Narayan, M.; Sirimulla, S., S…O and S…N Sulfur Bonding Interactions in Protein-Ligand Complexes: Empirical Considerations and Scoring Function. J Chem Inf Model 2016, 56, 2298–2309.

24. Beno, B. R.; Yeung, K. S.; Bartberger, M. D.; Pennington, L. D.; Meanwell, N. A., A Survey of the Role of Noncovalent Sulfur Interactions in Drug Design. J Med Chem 2015, 58, 4383–438.

25. Bannwarth, C.; Ehlert, S.; Grimme, S., GFN2-xTB-An Accurate and Broadly Parametrized Self-Consistent Tight-Binding Quantum Chemical Method with Multipole Electrostatics and Density-Dependent Dispersion Contributions. J Chem Theory Comput 2019, 15, 1652–1671.

26. Lang, P. T.; Holton, J. M.; Fraser, J. S.; Alber, T., Protein Structural Ensembles Are Revealed By Redefining X-Ray Electron Density Noise. Proc Natl Acad Sci U S A 2014, 111, 237–42.

27. Yao, S.; Moseley, H. N. B., A Chemical Interpretation Of Protein Electron Density Maps In The Worldwide Protein Data Bank. PLoS One 2020, 15, e0236894.

28. Musil, F.; Grisafi, A.; Bartok, A. P.; Ortner, C.; Csanyi, G.; Ceriotti, M., Physics-Inspired Structural Representations for Molecules and Materials. Chem Rev 2021, 121, 9759–9815.

29. Joosten, R. P.; Long, F.; Murshudov, G. N.; Perrakis, A., The PDB_REDO Server For Macromolecular Structure Model Optimization. IUCrJ 2014, 1, 213–20.

30. Liebschner, D.; Afonine, P. V.; Baker, M. L.; Bunkoczi, G.; Chen, V. B.; Croll, T. I.; Hintze, B.; Hung, L. W.; Jain, S.; McCoy, A. J.; Moriarty, N. W.; Oeffner, R. D.; Poon, B. K.; Prisant, M. G.; Read, R. J.; Richardson, J. S.; Richardson, D. C.; Sammito, M. D.; Sobolev, O. V.; Stockwell, D. H.; Terwilliger, T. C.; Urzhumtsev, A. G.; Videau, L. L.; Williams, C. J.; Adams, P. D., Macromolecular Structure Determination Using X-Rays, Neutrons And Electrons: Recent Developments In Phenix. Acta Crystallogr D Struct Biol 2019, 75, 861–877.

31. Liu, Z.; Su, M.; Han, L.; Liu, J.; Yang, Q.; Li, Y.; Wang, R., Forging the Basis for Developing Protein-Ligand Interaction Scoring Functions. Acc Chem Res 2017, 50, 302–309.

32. Schrodinger, LLC, In; 2015.

33. McKinney, W., Data Structures for Statistical Computing in Python. Proceedings of the 9th Python in Science Conference 2010, 445, 51–56.

34. Harris, C. R.; Millman, K. J.; van der Walt, S. J.; Gommers, R.; Virtanen, P.; Cournapeau, D.; Wieser, E.; Taylor, J.; Berg, S.; Smith, N. J.; Kern, R.; Picus, M.; Hoyer, S.; van Kerkwijk, M. H.; Brett, M.; Haldane, A.; del Río, J. F.; Wiebe, M.; Peterson, P.; Gérard-Marchant, P.; Sheppard, K.; Reddy, T.; Weckesser, W.; Abbasi, H.; Gohlke, C.; Oliphant, T. E., Array Programming With NumPy. Nature 2020, 585, 357–362.

35. Hunter, J. D., Matplotlib: A 2D Graphics Environment. Computing in Science & Engineering 2007, 9, 90–95.

36. InkscapeProject, Inkscape 0.92.5. 2020.

